# Revealing the Temporal Dynamics in Non-invasive Electrophysiological recordings with Topography-based Analyses

**DOI:** 10.1101/779546

**Authors:** Xuefei Wang, Hao Zhu, Xing Tian

**Author notes:** Corresponding author: Xing Tian, New York University Shanghai, 1555 Century Avenue, Room 1259, Shanghai, China 200122, (86-21) 20595201.

## Abstract

The fine temporal resolution of electroencephalography (EEG) makes it one of the most widely used non-invasive electrophysiological recording methods in cognitive neuroscience research. One of the common ways to explore the neural dynamics is to create event-related potentials (ERPs) by averaging trials, followed by the examination of the response magnitude at peak latencies. However, a complete profile of neural dynamics, including temporal indices of onset time, offset time, duration, and processing speed, is needed to investigate cognitive neural mechanisms. Based on the multivariate topographic analysis, we developed an analytical framework that included two methods to explore neural dynamics in ERPs. The first method separates continuous ERP waveforms into distinct components based on their topographic patterns. Crucial temporal indices such as the peak latency, onset and offset times can be automatically identified and indices about processing speed such as duration, rise, and fall speed can be derived. The second method scrutinizes the temporal dynamics of identified components by reducing the temporal variance among trials. The response peaks of signal trials are identified based on a target topographic template, and temporal-variance-free ERPs are obtained after aligning individual trials. This method quantifies the temporal variance as a new measure of cognitive noise, as well as increases both the accuracy of temporal dynamics estimation and the signal-to-noise ratio (SNR) of the ERP responses. The validity and reliability of these methods were tested with simulation as well as empirical datasets from an attention study and a semantic priming (N400) study. Together, we offer an analytical framework in a data-driven, bias-free manner to investigate neural dynamics in non-invasive scalp recordings. These methods are implemented in the Python-based open-source package *TTT* (Topography-based Temporal-analysis Toolbox).

## Introduction

The accessibility of non-invasive human scalp recordings, such as electroencephalography (EEG) makes it one of the most widely used tools in the cognitive neuroscience research as well as in the clinical and commercial settings (Biasiucci, Franceschiello, & Murray, 2019). Its fine temporal resolution at millisecond-level grants this technique enormous advantages to probe the underlying neural dynamics of various cognitive functions (Luck, 2014). However, non-invasive scalp recordings are usually noisy. To improve SNR, neural signals are averaged across trials to reveal relatively stable dynamics. With random noise being averaged out, the remaining waveform is called the *Event-Related Potential (ERP)*. One of the common ways to explore the neural dynamics in ERPs is to examine the neural response magnitude around certain peak latencies. Using this method, many important ERP components that reflect the processing dynamics of cognitive functions have been identified, such as N1, P2, P300, N400, P600 etc (Näätänen & Picton, 1987; Luck & Hillyard, 1994; Donchin & Coles, 1988; Kutas & Hillyard, 1980; Friederici, 2002).

The well-established ERP analysis utilizes only a small portion of the high-resolution temporal information from EEG recordings, namely the peak latencies. More temporal indices can provide a more complete picture of neural dynamics. For example, the *onset time* of a component can represent the initiation of a cognitive process. The difference between the onset time and the peak latency, the *rising duration*, reflects the accumulation speed of a cognitive process. Temporal information about ERP components can be used to answer many cognitive research questions. For example, faster neural activation or shorter duration was observed with the repetitive exposures of the same stimuli, which suggested neural facilitation as a potential mechanism to the behavioral priming (Grill-Spector, Henson, & Martin, 2006). The rich neural dynamics and its importance have been implicated in many other cognitive domains, such as aging (Dustman et al., 1990), learning (Rossion, Gauthier, Goffaux, Tarr, & Crommelinck, 2002), decision making (Nieuwenhuis, Aston-Jones, & Cohen, 2005), attention (Zopf, Giabbiconi, Gruber, & Müller, 2004), emotion (Streit, Wölwer, Brinkmeyer, Ihl, & Gaebel, 2000; Huang & Luo, 2006), memory (Haenschel, Vernon, Dwivedi, Gruzelier, & Baldeweg, 2005).

Despite the importance of the temporal information, the investigation of EEG dynamics is scarce. Estimating the temporal indices is constrained by the limitations of EEG recordings. First, signals from the same neural source may be picked up at different times by different sensors, because the distributed sensors have different distances from the source. It is difficult, if not impossible, for an observer outside the black box to accurately infer the temporal information of underlying neural processing from data of individual scalp sensors. Second, an ERP component may consist of multiple neural sources that overlap completely or partially in space and time. The complex spatial-temporal dynamics are hard to disentangle. Last, the common practice to boost the signal-to-noise (SNR) of EEG signals is to average multiple trials. However, the temporal features of signals vary across trials (e.g. may start earlier, process faster in some trials). This temporal variance is even greater in the components that are temporally further away from the onset of stimuli. When averaging trials, the actual temporal dynamics, and its parameters are smeared due to the temporal variance across trials.

Current common practices of temporal analyses are mostly based on the response magnitude of waveforms in individual sensors (Luck, 2014). For example, peak latency is defined as the time point when the voltage reaches the local maximum within a specific range. Onset and offset latencies are determined based on the relative response magnitude changes from a baseline measure. However, methods that rely on the magnitude in individual sensors are subject to the abovementioned problems -- the variation of dynamics across sensors, complex dynamics among sensors due to temporal and spatial overlapping of neural sources. Moreover, the practice of “picking sensor” is laborious, and may introduce subjective bias, and cannot represent the entire dynamics. Single sensor analysis is also subject to possible temporal variance induced by averaged single trials. Therefore, averaging and examining single sensor waveforms makes the investigation of temporal dynamics less optimal, if not misleading.

Recently, we developed an alternative ERP analysis method based on the topography (Tian & Huber, 2008; Tian, Poeppel, & Huber, 2011; Yang, Zhu, & Tian, 2018). A topography of ERP responses is the electric field distribution in all sensors over the scalp. It represents the configuration of underlying neural processes. We have developed multivariate analysis methods for testing psychological and neuroscience hypotheses by assessing the patterns of topographies between experimental conditions (Tian & Huber, 2008; Tian, Poeppel, & Huber, 2011) and implemented these methods in an open-source toolbox, EasyEEG (Yang, Zhu, & Tian, 2018). These topography-based multivariate analysis methods can distinguish changes in response magnitude vs. changes in neural source configuration, overcome the individual differences, avoid the pitfalls of using individual sensor data and provide an unbiased and complete assessment of underlying neural processes (Tian & Huber, 2008).

More importantly, the changes of topography over time can directly and unbiasedly reflect the neural dynamics. Therefore, in this study, we develop an analytical framework that includes two temporal analysis methods based on topography to probe the temporal dynamics of ERP components. Specifically, the first method (*Temporal evolution of components*) segments continuous waveforms into discrete ERP components and extracts a set of temporal information for each component. The second method (*Precise temporal estimation after reducing temporal variance across trials*) assesses the temporal variance across trials and provides an alignment approach to reduce variance for better evaluation of temporal parameters. These steps overcome the problems of variation across sensors and trials and offer an analytical framework in a data-driven, bias-free, automatic manner to investigate neural dynamics in non-invasive human scalp recordings. These methods have been implemented in an open-source toolbox, the Topography-based Temporal-analysis Toolbox (*TTT*, https://github.com/TTT-EEG/TTT-EEG). It also offers an interface to effectively link with our ERP data analysis software EasyEEG (Yang, Zhu, & Tian, 2018, https://github.com/ray306/EasyEEG) and python-based MNE software (Gramfort et al. 2013, https://github.com/mne-tools/mne-python).

In the following sections, we first provide an introduction of each method, followed by a test case using simulated data with predetermined characteristics to illustrate its functionality. Lastly, we apply each method on empirical data to further validate the method in a real experimental context.

## Temporal evolution of components

A cascade of cognitive processes unfolds over time to effectuate a behavior. ERP waveforms manifest these cognitive processes as consecutive components. It’s important to separate each component in the temporal space and evaluate its starting and ending points. This separation helps to better identify and quantify the dynamics of cognitive processes, and to facilitate later analysis. Therefore, we first propose a method to separate continuous waveforms into distinct components and to extract a set of temporal indices (onset time, peak latency and offset time) for each component. This method assumes that a topographic pattern remains consistent throughout the period of an ERP component, and changes of topographic patterns indicate the transition between components. We used both datasets from artificial simulation and real experiments to test the validity of this assumption and method. Both results strongly supported our assumption and method.

### Method

An ERP component that reflects specific cognitive processes usually takes time to unfold. Throughout the time course, the response magnitude waxes and wanes. Whereas, the response pattern in the sensor space (topography) remains relatively consistent because the configuration of the underlying neural sources remains the same. Therefore, identifying the transition between topographies can separate components in time, and provides temporal boundaries (the onset and offset time points). Whereas, the time point when response magnitude reaches maximum yields the peak latency of that ERP component. The complete set of temporal indices for an ERP component (onset, peak, and offset time) provides the necessary information to define the evolution of that component.

The intra-component topographies exhibit a high degree of pattern similarity while inter-component topographies are dissimilar. Therefore, we use the degree of similarity between topographies to identify the temporal boundaries (onset and offset times) between components. Cosine distance measure (Manning, Raghavan, & Schütze, 2010; Tian & Huber, 2008) quantifies the degree of similarity between topographies. Specifically, a topography is mathematically a high dimensional vector, where the number of dimensions equals the number of sensors. Cosine value of the angle between two vectors represents the degree of similarity between two topographies ***A***, *B* -- the larger the cosine value is, the more similar the topographies are (Tian, Poeppel, & Huber, 2011):

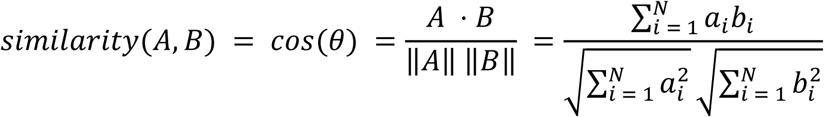

The cosine value is independent of response magnitude and only represents the response pattern similarity. It is the key feature that enables the detection of the transition of topographies between components.

An epoch with *N* time points is used as an example for demonstration (*N* = *f* * *T*, where *f* is the sampling rate and *T* is the duration). A topography is presented at each time point (Fig. 1A). We constructed an N-by-N similarity matrix (Fig. 1B) by comparing each pair of topographies. The (i^th^, j^th^) element of the matrix denotes the degree of similarity between the i^th^ and j^th^ topographies. The similarity matrix is symmetrical along the principal diagonal line. All cosine values are in the range of [-1, 1].

**Figure 1.**
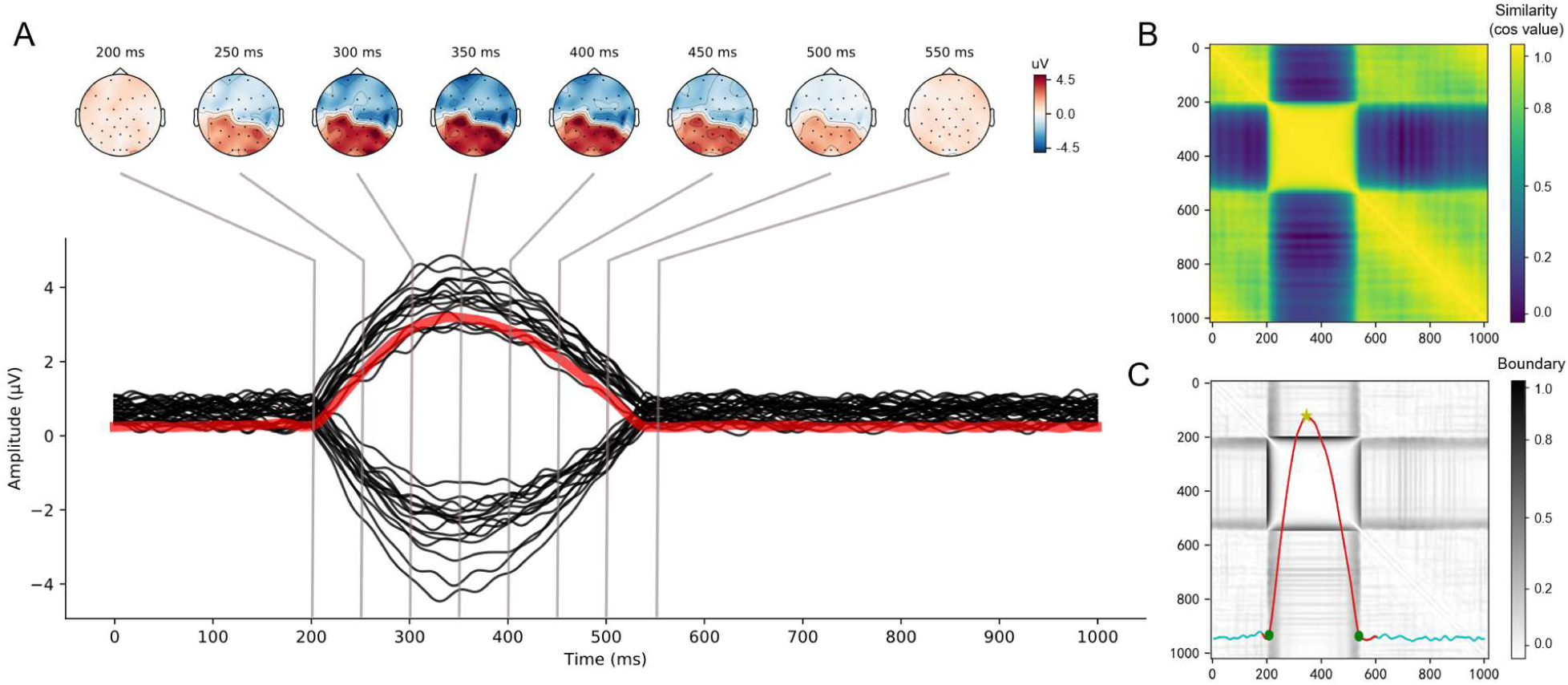
Simulation results of the first method (Temporal evolution of components). (A) The simulated data. The simulated waveform responses were generated using sinusoidal waves with additional Gaussian noise. Black lines in the lower panel represent the raw data of 32 sensors. The red line represents the global field power (GFP), the geometric mean of responses across all sensors. Topographies at selected latencies are depicted in the upper panel, showing the temporal evolution of the component. (B) The similarity matrix. Cosine values are obtained by comparing each pair of topographies (1000 × 1000). The (i^th^, j^th^) element of the matrix denotes the degree of pattern similarity between the i^th^ and j^th^ topographies. Principal diagonal line is the auto-correlations of each topography, therefore the values are all ones. High-correlation (green-yellow) clusters along the principal diagonal line reveal the component. (C) Gradient matrix with superimposed GFP waveform. Dark ridges in the gradient matrix represent the cluster boundaries. The GFP waveform is superimposed on the gradient matrix with the red portion represents the pre-selected period of interest (POI) for the detection of a component. The green dots on the GFP label the detected onset and offset time points, corresponding to the ridges in the gradient matrix. The yellow star represents the detected peak latency at the maximum amplitude of the GFP waveform.

With the similarity matrix, we identify square-shaped high-similarity clusters in the matrix to identify each component. The boundaries of a cluster are the onset and offset time points of an ERP component. To capture the clusters automatically, we use an edge detection operation with Scharr kernel (Scharr, 2000; Jähne, Haussecker, & Geissler, 1999) on the similarity matrix. We define the Scharr kernel as *S*, the similarity matrix as *K*. The Scharr operator *S* is defined as:

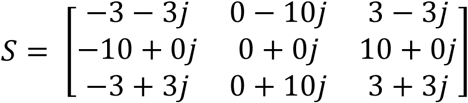

The edge detection operation is implemented as a 2-D convolution. By applying the convolution to the *K* and *S*, we produce a gradient matrix denoted as *G* (Fig. 1C):

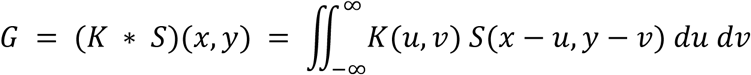

The ridges in the gradient matrix are the boundaries of the clusters. Therefore, we can obtain the precise onset and offset time points of an ERP component.

Based on the detected onset and offset times, we can derive more indices for the measurement of ERP components’ temporal dynamics. The first index is the *duration* of an ERP component. The duration represents the total activation time of an ERP component and reflects the efficiency of performing underlying cognitive processes. The second index is the *rise speed* (RS) and *fall speed* (FS). These two indices are the changing speed of response magnitude from onset to peak, and peak to offset, respectively. They capture how fast neural activity accumulates and decays, which is asymmetric in some cases. We suggest that these indices reflect the intrinsic temporal properties of ERP components. Hence, they are important measures to investigate the dynamics of cognitive processes (Ng, Tobin, & Penney, 2011).

### Simulation

We generated a simulation dataset to test the method. Each topography had 32 sensors with a standard 10-20 montage. The sampling rate of the dataset was 1000 Hz. The total duration of the dataset was 1000 milliseconds. One ERP component was simulated by concatenating a rising half and a falling half based on sinusoidal waveforms (rising duration: 140ms, falling duration: 200ms). Each sensor shared the same phase, but differed in amplitudes (normal distribution, mean = 3.00μV, std = 0.60μV, max = 4.59μV, min = 1.83μV). The onset, peak, offset time points were set to 200ms, 340ms, 540ms, respectively. We also added a Gaussian noise (frequency range: 0.1-30 Hz, SNR = 2.5) to the signal. The simulated dataset is presented as a waveform plot with the global field power (GFP) superimposed (Fig. 1A). Topographies at selected time points are shown above the GFP waveform to demonstrate the temporal evolution of the component. The cosine similarity between a pair of topographies at each time point was calculated and presented as a 1000-by-1000 similarity matrix (Fig. 1B).

To automatically and precisely capture the onset and offset time points, we used the Scharr kernel to generate a gradient matrix (Fig. 1C). The ridges (dark lines in Fig. 1C) in the gradient matrix plot denotes the temporal boundaries of the component. The red portion of the GFP waveform denotes the preselected window that contains the component. The green dots denote the automatically detected onset and offset time points and yellow star denotes the peak latency. Our method successfully detected the onset and offset latencies of the pre-constructed components (onset: 203ms, peak latency: 341ms, offset: 536ms, duration: 333ms, rise speed: 20.81μV/s, fall speed: 15.06μV/s).

### Application

We used an empirical dataset from a recent ERP study (Zhang, Tao, & Zhao, 2019) to further test our method. This study investigated how perceptual separation and auditory spatial attention interact with each other to facilitate speech perception in a noisy environment. Latency differences in N1 and P2 components under different conditions were reported. We applied our method on this dataset to further test the validity and efficiency of our method.

First, our method identified N1 peak latency based on the individual GFP waveform for each condition. Then we performed a repeated-measures two-way ANOVA with factors of *perceptual location* (2 levels) and *direction of attention* (2 levels) on the N1 peak latency. The main effect of *perceptual location* was significant (*F*(1,18) = 5.122, *p* = 0.036). Neither *direction of attention* (*F*(1,18) = 2.862, *p* = 0.108) nor the interaction (*F*(1,18) = 0.180, *p* = 0.677) reached significance. Our results on N1 latency were consistent with the original findings (Zhang, Tao, & Zhao, 2019). Our method could not reliably detect the peak of the P2 component because there was no clear peak in the GFP waveform (see Fig. 2A top-right as an example). It was also consistent with the speculation in the original paper that P2 and later P300 component might temporally overlap. Our automatic method successfully replicated the findings in an empirical ERP study, suggesting the validity and efficiency of our method when applied in a practical setting.

**Figure 2:**
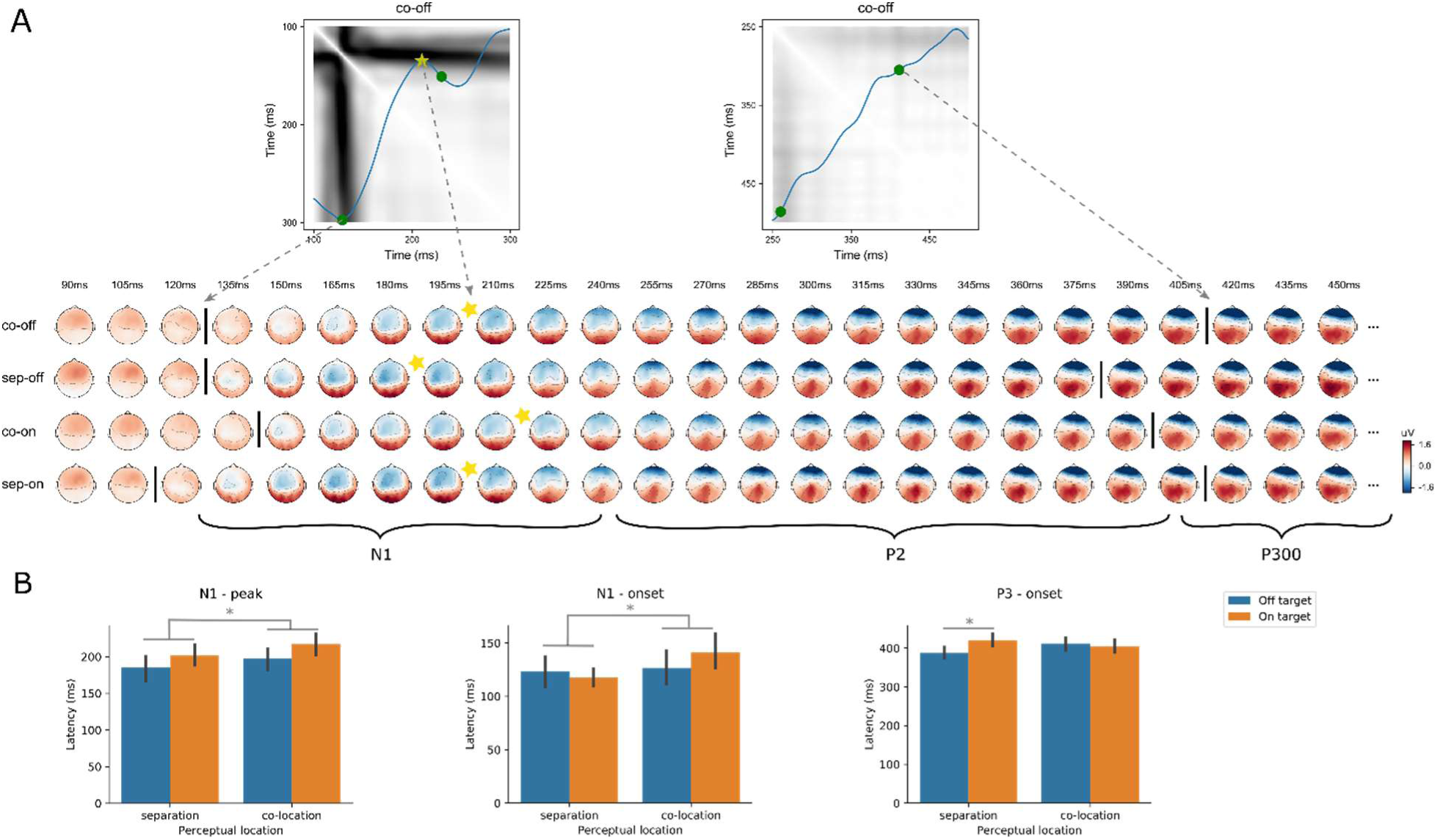
Results of the first method (*Temporal evolution of components*) on an auditory spatial attention experiment (Zhang, Tao, & Zhao, 2019). (A) Extracted temporal indices from the empirical dataset. Lower panel, topographies across time for each experimental condition. Upper panel, gradient matrices with GFP and detected temporal indices (onset, offset, and peak latency, similar to Fig. 1C). Abbreviation: co: perceptual co-location, *sep*: perceptual separation, *on*: on-target, *off*: off-target. (B) Statistical test for the N1 peak latency, N1 onset, and P300 onset. The main effect of perceptual location was significant for the N1 peak latency, as well as the N1 onset. The interaction was significant for P300 onset latency.

Moreover, we supplemented the original findings with extra temporal indices (N1 onset, offset; P2 onset, P300 onset -- the temporal boundary between P300 and P2) that were automatically identified by our method. Similar repeated-measures two-way ANOVA was performed on each index. First, for the N1 onset, the main effect of *perceptual location* was significant (*F*(1,18) = 4.889, *p* = 0.040). But *direction of attention* was not significant (*F*(1,18) = 0.353, *p* = 0.560), neither was the interaction (*F*(1,18) = 3.468, *p* = 0.079). These results of the N1 onset time point differences suggest that the perceptual location modulates the initiation timing of the very first auditory ERP response. The originally observed difference in the N1 peak latency was likely caused by the early initiation of neural responses in the perceptual co-location condition. Second, for the P300 onset, the main effects were not significant (for *perceptual location, F*(1,18) = 0.309, *p* = 0.585, for *direction of attention, F*(1,18) = 3.145, *p* = 0.093). However, a significant interaction was observed (*F*(1,18) = 11.708, *p* = 0.003). Further t-test showed that the onset time of P300 component in the *on-target attention* condition was later than that in the *off-target* condition when the perceptual locations were separated (*t*(1,18) = 2.404, *p* = 0.027). Neither N1 offset or P2 onset showed significant differences among conditions. These results suggest that the faster transit from P2 to P300 possibly reflects the processing speed of cognitive mechanism between perceptual location and direction of attention in a “cocktail party” environment. Using our new method, we obtained additional temporal dynamics results that were crucial evidence to test potential neural mechanisms. These results suggest that our automatic topography-based temporal segmentation method can provide a complete set of temporal measures to investigate various aspects of cognitive neuroscience theories.

## Precise temporal estimation after reducing temporal variance across trials

Temporal indices can be captured by the first topography-based method. It overcomes the problems of variance across sensors as well as temporal and spatial overlapping of neural sources in the sensor space. However, they are still based on averaging of trials, and therefore are subject to temporal variance across trials. For example, if response peaks in individual trials have large temporal variability and are not temporally aligned, the duration of an averaged ERP component would be longer than it truly is. Moreover, because of the temporal unalignment among individual trials, the response magnitude could be averaged out and hence smaller than the actual peak response. In most cases, the temporal variance among trials is what researchers should be cautious to avoid. Whereas in other cases, the temporal variance among trials is the measure of interest and implies possible underlying cognitive mechanisms. For example, one study showed that children with ASD had more variance in their P1 latency to Gabor patches than the control group, lending support to the theory of increased neural noise in ASD (Milne, 2011). Therefore, it’s necessary to quantify the temporal variance among intra-subject single trials, followed by either removal the temporal variance to obtain a more precise estimation of temporal indices, or analysis the temporal variance to investigate neural and cognitive theories.

Here, we propose a method to find peaks in single trials based on a topographic template. This method assesses the temporal variance across trials and obtains temporal-variance-free ERPs by aligning trials. This method can increase the accuracy of temporal dynamics estimation. The realignment can also increase the SNR of the averaged ERP responses. Furthermore, this method takes the temporal variance across trials into account and offers a novel angle to explore the ERP components -- whether the experimental manipulations change the temporal dynamics of a component or change the variance across trials.

### Method

Peak latency of a component in an ERP can be identified using the first method. The topography at the peak latency is selected as a template, denoted by *T* (Fig. 3A). This selected template topography is the stable and accurate neural representation of the component. Topography can be viewed as a vector in a high dimensional space, and the vector of template topography provides the exact direction of the ERP component.

**Figure 3:**
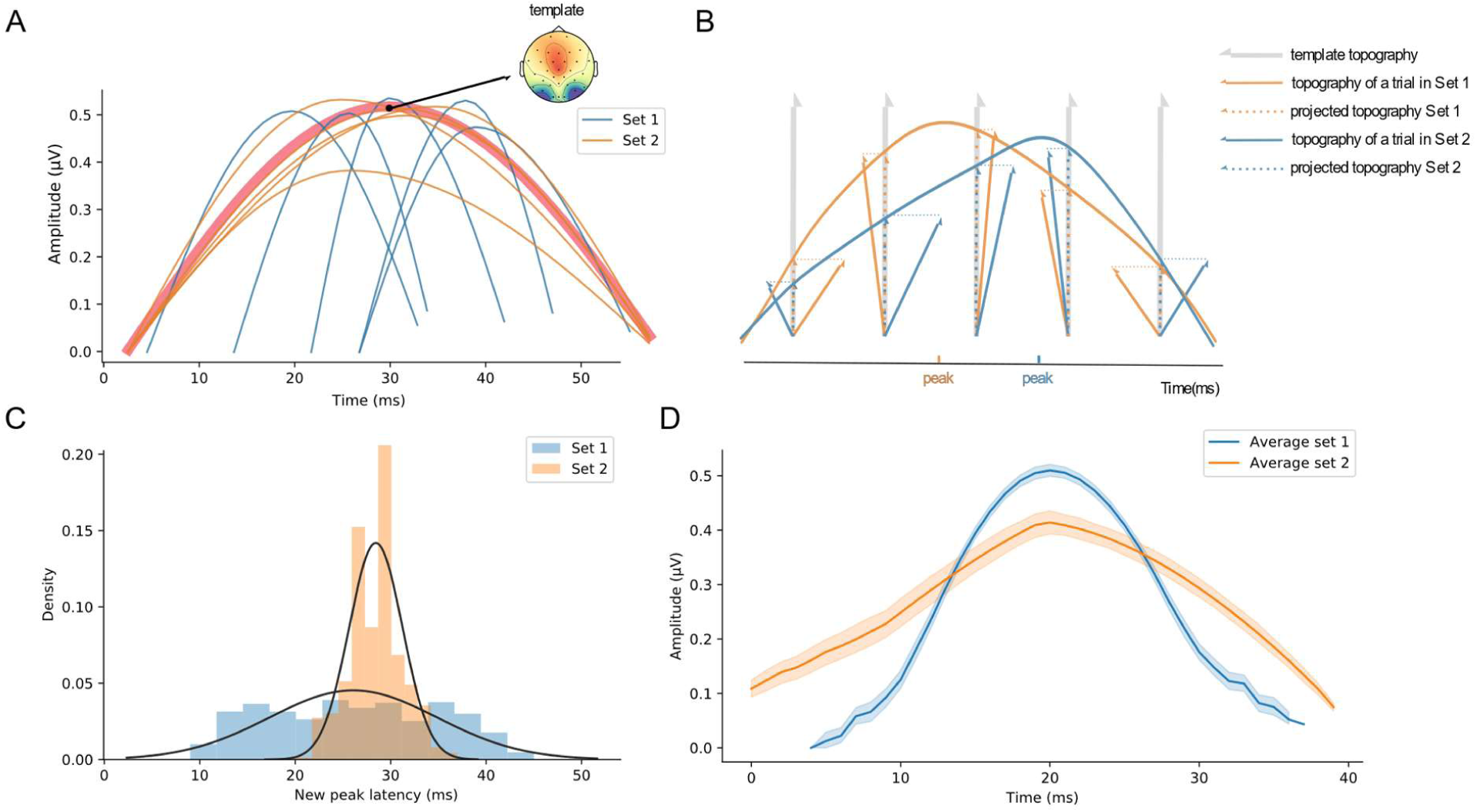
Simulation results of the second method (*Precise temporal estimation after reducing temporal variance across trials*). A) Two types of simulated trials yield the same averaged waveform. The waveforms represent the GFP of simulated data. The first set of trials are depicted in blue. These trials have a short duration and high temporal variance of peak latencies among trials. The second set of trials is illustrated in orange. These trials have a long duration and low temporal variance of peak latencies among trials. The average of both sets of trials yields the same ERP responses (bold red waveform). A template topography *T* is selected at the peak latency of the averaged responses. B) Schematic plot of projection for identifying peak latency in single trials. The gray arrow represents the vector of template topography, whereas the blue and orange arrows represent the topographies of a single trial across time from the first and second sets. Each topography at a time point is projected onto the template topography *T*. The length of the parallel part represents the noise-free response magnitude. The projected values form new waveforms and the new peak latency is identified. C) Distributions of individual trial latencies. Histograms of new peak latencies identified in B) are depicted in corresponding colors (blue for the first set trials and orange for the second set). The normal distribution function is fitted to both histograms. The first set of trials results in a flat and wide distribution (blue, *μ* = 26.09, *σ* = 8.80). Whereas the second set of trials yields a tall and sharp distribution (orange, *μ* = 28.45, *σ* = 2.81). D) Averaged ERP responses after aligning trials at new peak latencies. The second set of trials yield an average response that is similar to the original one in A). Whereas the average response of the first set of trials is shorter in duration and higher in amplitude compared with the original. color shaded areas represent +/- standard error of the mean (SEM).

For every trial, the topography *t*^*i*^, at each time point *i* is projected onto the vector of template topography *T* (Fig. 3B). In this way, each topography vector is decomposed into two orthogonal parts, one parallel to the template, 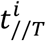, and the other perpendicular to it, 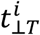:

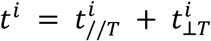

The perpendicular part that is orthogonal to the direction of template topography, is irrelevant to the representation of interest. Whereas, the strength of the parallel part reflects the activity along the direction of the template vector. By only keeping the parallel part, we can measure the noise-free response magnitude as the projection length *c*^*i*^:

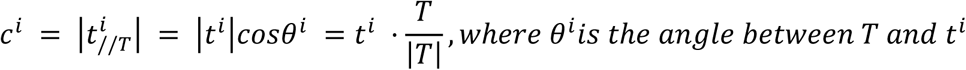

The projection calculation is carried out for each time point during the duration of the component. The projected values as a function of time form a new waveform for each trial. The time point when the single-trial projection length *c*^*i*^ reaches the maximum is identified as the new peak latency for each trial and denoted as *k*:

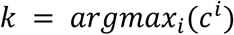

A distribution of single-trial peak latencies relative to the template latency can be obtained (Fig. 3C). To estimate the temporal variance of individual trials, this distribution is fitted with a probability density function. The temporal variance can be estimated in the fitted distribution. After identifying individual trial peak latencies, we can align trials according to the individual peak latencies and obtain the ERP by averaging the aligned trials (Fig. 3D). This temporal alignment across trials makes the peak of a given component time-locked and hence increases the SNR and reveals the undistorted true waveform.

### Simulation

To test this method, we generated two sets of trials based on a sinusoidal waveform (Fig. 3A). Each set of trials has different single trial patterns but results in identical averaged waveforms (bold red waveform in Fig. 3A). The trials in the first set (trials in blue in Fig. 3A) have a shorter duration and higher varied peak latencies than the trials in the second set (trials in orange in Fig. 3A). Specifically, for the trials in the first set, the duration for each trial is sampled from a uniform distribution *x* ∼ *U*(20, 30), and its onset is from *x* ∼ *U*(0, 30). The peak occurs around the middle of the duration with a jitter from a uniform distribution of *x* ∼ *U*(0.4, 0.6). The amplitude is set to be randomly sampled from *x* ∼ *U*(0.4, 0.6). For the second set, the onset (0 ms) and duration (55ms) of all trials are set to be the same, while peak occurs at different time points in around the middle point jittering from a uniform distribution of *x* ∼ *U*(0.45, 0.65). The amplitude is also sampled from *x* ∼ *U*(0.4, 0.6).

The single-trial latencies in each set are identified using this method after projecting topographies of individual trials onto the template (Fig. 3B). The identified peak latencies in the first set of trials yield a flat and wide distribution (Fig. 3C, blue). A normal probability density function is fitted against the distribution (*μ* = 26.09, *σ* = 8.80). Whereas the identified peak latencies in the second set of trials reveal a tall and sharp distribution (Fig. 3C, orange). The fitted parameters are *μ* = 28.45, *σ* = 2.81. These results suggest more temporal variance among trials in the first set than the second set, which is consistent with the pre-determined settings. Moreover, after aligning to the identified new peaks, the new average ERP of the first set (Fig. 3D, blue) is narrower and taller than the original averaged ERP waveform (Fig. 3A, red) and resembles more closely to the processing dynamics of individual trials (Fig. 3A, blue). This suggests improved SNR and better estimation of temporal indices for the first set of trials that have more temporal variance.

### Application

We used an unpublished empirical dataset from a semantic priming study to further demonstrate the effectiveness of this method. Participants were asked to read letter strings that could be words or nonwords. They performed a lexical decision task. The early visual response of N1 and the semantic-related response of N400 components (Kutas & Federmeier, 2011) are identified using the first method (Fig. 4A, N1 latency: 86ms, N400 latency: 412ms). The N1 component has a sharp peak, with a short duration, presumably because the transit lower-level visual processing is temporally close to and well time-locked to the onset of stimuli. Whereas the N400 component exhibits a wide, multiple-peak profile. The wide waveform can be explained by several hypotheses. The first is that there is one cognitive process (component) that have a long duration and such a long process is relatively consistent among individual trials, similar to the simulated Set 2 orange trials in Fig. 3A. The second possible scenario is that there is only one cognitive process (component), but the process is short and the peak latencies in individual trials vary as the semantic-related process is further away from the onset of stimuli, similar to the simulated Set 1 blue trials in Fig. 3A. The last one is that it may contain multiple cognitive neural processes that unfold over time with temporal overlaps. However, the topographies have similar patterns during the duration of N400. Therefore, the last hypothesis is unlikely. The remaining two hypotheses compete in terms of the true cause of the observed long duration in an ERP component -- whether the observed long-duration N400 component is because of the underlying temporal dynamics, or the temporal variance across trials.

**Figure 4:**
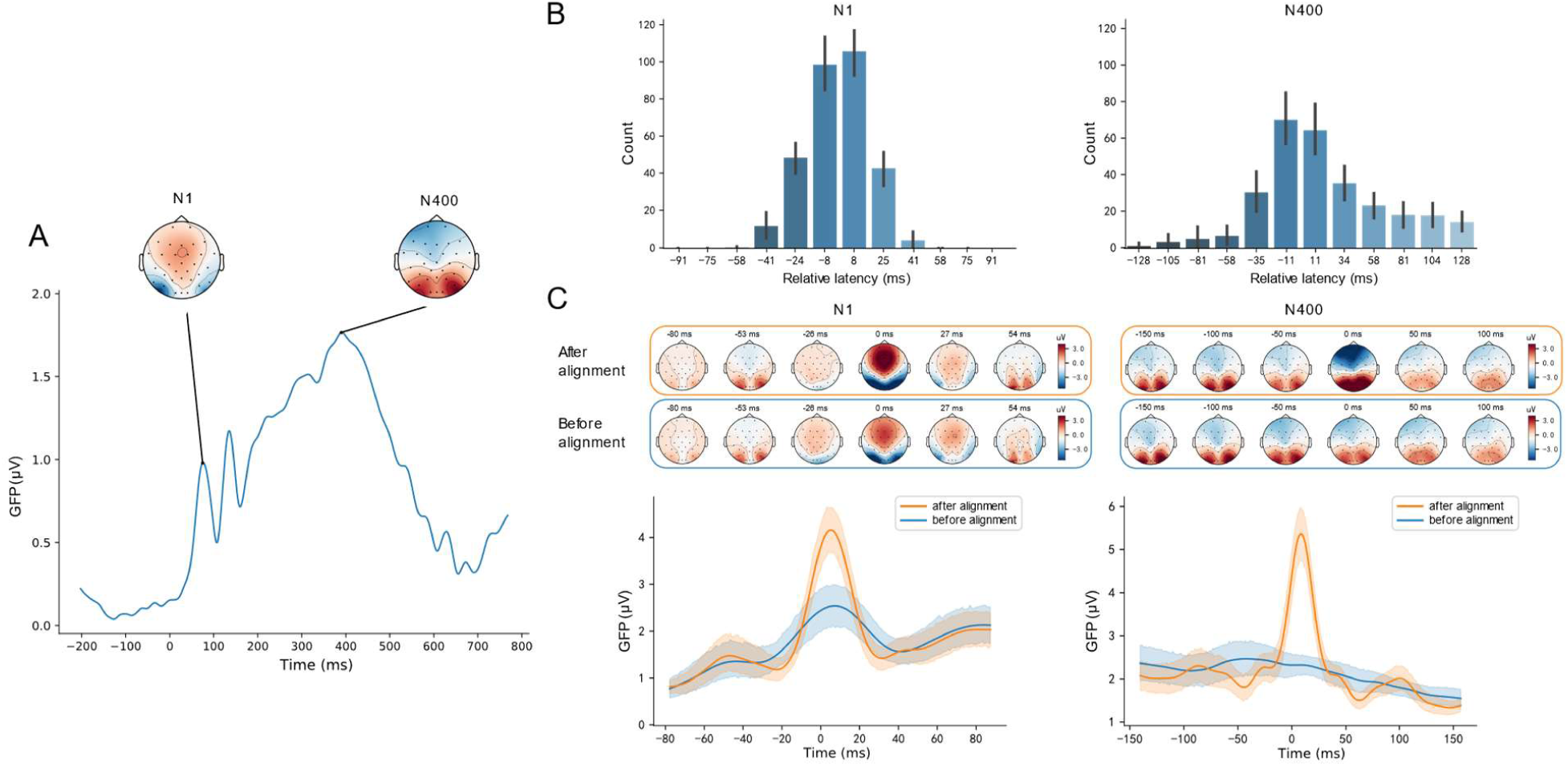
Results of the second method (*Precise temporal estimation after reducing temporal variance across trials*) on a semantic priming experiment. A) Original averaged GFP from a semantic priming study (unpublished data). The topographies of N1 and N400 components are shown. B) Distribution of single-trial peak latencies for N1 and N400 (Similar to Fig.3C). C) Topography and GFP before and after alignment for N1 and N400. The peaks of both components become sharper. The SNR of N400 was better improved compared with N1. The estimated duration of N400 was shorter than the original data in A). The shaded areas represent +- standard error of the mean (SEM).

We used the second method to probe the temporal characteristics of single trials that were averaged and resulted in the N400 and N1 components. Peak latencies of the N1 and N400 components were identified for every trial and were compared to the averaged peak latency. The relative latencies (single trial peak latency minus averaged peak latency) are plotted as histograms (Fig. 4B). A tall, narrow distribution pattern is shown for the N1 single trial peak latencies. Whereas a flat, wide distribution for N400. These results indicate that the temporal variance of peak latencies across individual trials are much larger in N400 than in N1. The long duration and multiple peaks in N400 are presumably caused by the temporal variance in single trials. Consider the temporal variance in the N1 as a null hypothesis (the baseline level of temporal variance across trial caused by the systematic neural noise), the larger temporal variance in the N400 suggest additional contribution from the cognitive-level noise.

We further aligned the trials to boost the SNR. The average responses of both N1 and N400 become sharper after alignment (Figure. 4C), and the effect is more prominent for N400 than N1. The topography intensity is also enhanced. To quantify the SNR improvement, we calculated the GFP peak amplitude ratio between the aligned-average and original waveforms. The SNRs for N1 and N400 were 1.76 and 2.29, respectively.

After aligning the peak latencies of individual trials, only one peak was observed in the N400 component (Fig. 4C). Moreover, the duration of the aligned-average N400 (Fig. 4C) was much shorter than the original one (Fig. 4A). Together with the observation of large temporal variance across trials for the N400 (Fig. 4B), these results support the hypothesis that one short cognitive process mediates the N400 and the apparent long duration is caused by large temporal variance across trials. The temporal profile of semantic processing is better estimated in the peak-aligned average response.

## Discussion

We developed an analytical framework that includes two methods to investigate the temporal characteristics of ERPs. Both methods are based on topography, a multivariate representation of neural responses in non-invasive scalp recordings. The first method reveals the temporal evolution of ERP components by identifying crucial temporal indices (onset, offset, peak latency, duration, rise speed, fall speed). It provides a complete temporal profile of each ERP component. The second method estimates and reduces the temporal variance across trials within each ERP component. The second method yields a more precise estimation of temporal indices and higher SNR. Estimation of temporal variance with this method also establishes a new way of single-trial analysis to investigate theories regarding temporal variation and cognition. These two methods collaboratively provide a quantitative framework to investigate neural dynamics in non-invasive scalp recordings.

Our new methods target the temporal aspects of ERP. Previous methods use the temporal index, mostly peak latency, as a factor to constrain the main measure of response magnitude. The two new methods proposed in this study provide quantitative approaches to measure a complete set of temporal indices, including onset, offset, peak latency, duration, rise speed, fall speed. This complete temporal profile can reflect all aspects of neural dynamics, including initiation time, processing duration, accumulation and decay speed, which opens a new dimension in the analysis of non-invasive scalp recordings and pave a new way to test temporal aspects of cognitive neuroscience theories.

The proposed two methods are based on the analysis of topography. The topography-based analysis offers advantages over individual-sensor based analysis in several aspects. First, while the waveform of individual sensors is highly dependent on the choice of reference, the topography is reference-free (Murray, Brunet, & Michel, 2008) or can be used to achieve reference-free (Yao, 2001). Second, topography-based analysis utilizes all sensors, gives an aggregate result. Hence it is free from sensor picking bias, avoids the multi-comparison problem, and is robust to cross-sensor differences. Third, as a multivariate analysis, topography offers sufficient information to identify the ERP components in a data-driven, automatic way.

That these methods are based on topography also results in a novel segmentation approach. These methods are based on the assumption that the response patterns of ERP components are consistent throughout the period of activation. This assumption holds in most empirical data. Therefore, temporal continuity is enforced in our methods. In comparison, clustering methods do not guarantee the temporal continuity and sometimes has to add this constraint by additional conditions (Pascual-Marqui, Michel, & Lehmann, 1995). Moreover, because our method is based on matrices, the most significant boundary is determined by the topography that maximizes all distance with all other topographies. It makes the analysis more resistant to noise and hence yields more robust results and may increase internal consistency (Thigpen, Kappenman, & Keil, 2017).

These advantages of our methods have been demonstrated in the applications when we applied our methods on empirical data. First, additional temporal parameters provided insights from a new perspective to understand neural mechanisms. For example, our first method successfully identified the onset time of N1, in addition to the peak latency in an auditory attention experiment (Zhang, Tao, & Zhao, 2019). The results of peak latency replicated the original findings. More importantly, we found that the significant differences among conditions can be obtained early in the onset time of N1. These results suggest that the actual differences in temporal processing may start in the initiation stage.

Second, dissecting the temporal dynamics from different angles offers new perspectives to investigate neural and cognitive theories. For example, our second method estimated the temporal variance among trials. This estimation provided evidence suggesting that the observed long duration of N400 in a semantic priming experiment was induced by the variation of semantic processing speed across trials. Moreover, after reducing the temporal variance by aligning individual trials, the more precise estimations of processing duration and response magnitude for semantic retrieval were obtained. These results suggest that our methods can distinguish whether the observed neural dynamics is caused by cognitive noise across trials or the processing speed of a cognitive function. Such precise estimation separates the contribution of two sources on the observed neural dynamics in ERPs, which offers a new perspective of temporal variance to test neural and cognitive theories (e.g. Milne, 2011).

Third, the automatic detection of temporal indices based on topography overcomes many disadvantages in a manual selection based on data in single sensors. In addition to obvious advantages such as avoiding subjective bias, free from variation among sensors, and release from tedious, repetitive, and error-prone laborious manual procedures, the most important advantage is the boost of accuracy and reliability on the identification of temporal indices. For example, in the original auditory attention study (Zhang, Tao, & Zhao, 2019), the peak latency of the late perceptual responses was hard to find, presumably because the temporal overlaps between the P2 component and the following P300 component. The successive activation of two components resulted in the monotonous increase of response magnitude, and hence the local maximum -- the response peak was absent (Fig. 2A). Using the topographic pattern as an additional feature, our method successfully identified temporal boundary between P2 and P300 and obtained the onset time of P300. The onset time of P300 suggests the interaction between auditory attention and target location, which is hard to obtain using the response magnitude in sensors.

The two new methods aim to obtain a complete set of temporal indices for investigating the neural dynamics in ERP responses. Each method weights on different aspects of temporal estimation. We provide a recommended procedure for using these methods. First, determine the component(s) of interest. Use the first method to estimate the onset time, peak latency and offset time of the component(s) in each condition for each participant. Other temporal indices, such as rise and fall speed can be derived from these three primary estimations. After obtaining these temporal indices, standard inference statistics can be applied to test specific hypotheses. Second, use the second method on the single trials of each condition for each participant. The distribution of temporal indices can be obtained for the component(s) of interest. Temporal variance can be estimated for each condition in each participant, and standard inference statistics can be applied to compare the temporal variance changes by experimental manipulations. Moreover, the temporal variance of early perceptual responses (e.g. N1) can be used as a baseline to quantify the variance of later components. Furthermore, the new ERP responses can be obtained by aligning signal trials. The SNR-boosted response magnitude and more precise temporal indices can be used to further test the research questions. We have implemented the methods in the *TTT* package, an open-source toolbox in Python. The interface of importing and exporting ERP data with other Python-based packages are provided, including MNE-Python (Gramfort et al. 2013; Gramfort et al. 2014) and EasyEEG (Yang, Zhu, & Tian, 2018). Sample code snippets for illustrating the major part of the recommended workflow is included in the supplementary materials. More analysis scripts are available at https://github.com/TTT-EEG/TTT-EEG.

## Conclusion

We proposed an analytical framework that includes two new methods to probe the temporal characteristics of ERP responses. These methods provide automatic approaches to extract crucial temporal information. Moreover, temporal variance across trials can be estimated. These two methods fully exploit the fine temporal resolution of non-invasive scalp recordings, which leads to a new direction on data analysis to explore neural dynamics and creates a new dimension to test neural and cognitive theories.

## Acknowledgments

We thank Changxin Zhang for providing EEG data in the auditory attention experiment. This study was a portion of X.W’s project that was supported by the Summer Undergraduate Research Program (SURP) at the NYU-ECNU Institute of Brain and Cognitive Science at NYU Shanghai. This study was supported by the National Natural Science Foundation of China 31871131, Major Program of Science and Technology Commission of Shanghai Municipality (STCSM) 17JC1404104, and the Program of Introducing Talents of Discipline to Universities, Base B16018.

